# A novel cell communication method reveals that grik4 and gabrd may be critical for inducing death in RGNNV-infected groupers

**DOI:** 10.1101/2024.02.27.582406

**Authors:** Tengfei He, Yepin Yu

## Abstract

To investigate the infectious mechanism of RGNNV, we adopted multi-omics methods to study the precise cellular interactions. We combined proteomic, bulk-RNA seq and sc-RNA seq to search for the secret of RGNNV’s influence on the nervous system in grouper. Besides, we created a sc-RNA seq workflow for cell communication analysis that can be applied to those non-model organisms with a reference for the first time, which usually been done by comparing homologous genes in humans or mice in the past (Cheng, Chen et al. 2023), but we use stringdb database to predict interactions at the whole proteomic level (Szklarczyk, Kirsch et al. 2023), and we also build a R package of this procedure to help achieving this goal in other non-model organisms. The results shows that grik4 and gabrd might be the direct causes of death in RGNNV-infected groupers. We also present a mechanistic picture of RGNNV attacking the nervous system of grouper and causing nerve necrosis.

---

The multi-omics data were derived from published literature. The proteomic and bulk-RNA seq data were published by (Ge, Lin et al. 2020), can be downloaded from https://www.ebi.ac.uk/pride/archive/projects/PXD012763 (Proteomic), https://www.ncbi.nlm.nih.gov/geo/query/acc.cgi?acc=GSE126620 (Bulk-RNA seq), respectively. And the sc-RNA seq data was published by (Wang, Peng et al. 2021), can be downloaded from https://www.ncbi.nlm.nih.gov/bioproject/PRJNA773515/ (Sc-RNA seq). And the reference genome of grouper we used is from *Epinephelus lanceolatus* can be downloaded at https://www.ncbi.nlm.nih.gov/datasets/genome/GCF_005281545.1/ (Reference genome).

The former experiment set a control group in vivo with 3 replicates, and two infection groups, which 3 replicates were moribund, and other 3 replicates were alive before execution. The later set two groups in vivo, the control group was injected with PBS, and the infection group with RGNNV suspension injection. Fig 1 A shows the schematic of experimental design, and details were described in the published materials mentioned before.

**Fig. 1.**
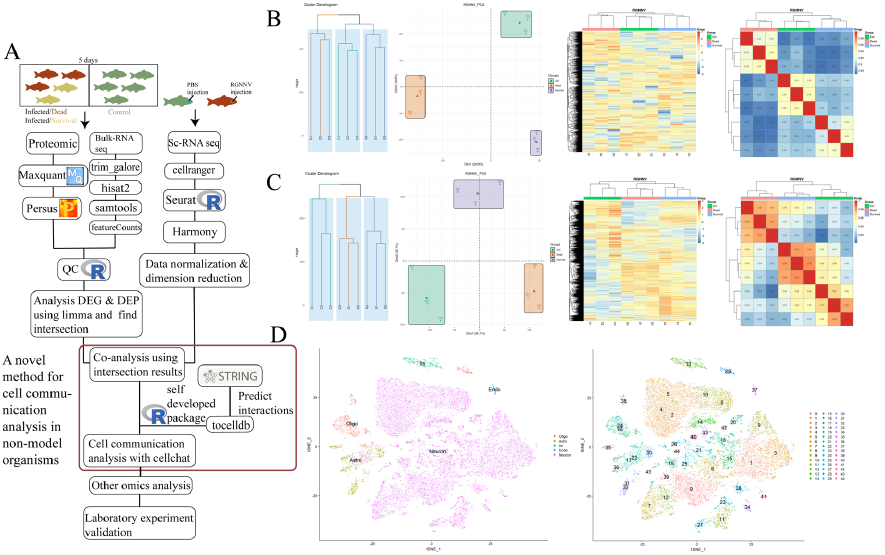
Data analysis overview. A. Sampling schematic and multi-omics analysis workflow. B. Proteome QC plot. C. Bulk-RNA seq QC plot. D. Sc-RNA seq tSNE plot.

## Data pre-processing

### 1. Proteomic data

We use Maxquant software to quantification the RAW data generated by label-free LC-MS, Label-free quantification set to ‘LFQ’, the proteomic sequences were downloaded from NCBI(GCF_005281545.1), and other parameters with default (Cox and Mann 2008, Tyanova, Temu et al. 2016). Then we filtered the data with Persus software, and 1927 protein was successfully quantified after filtration (Tyanova, Temu et al. 2016). The log2 transformed data was used for quality control checks in R. Fig 1 B shows the data within the same group were highly consistent, and there were obvious differences between the control and treatment groups.

### 2. Bulk-RNA seq

We use trim_galore to filter RNA sequencing data with q25 qualities and cut adaptor with length 3 first (Simons 2010, Martin 2011). Then hisat2 and samtools was used for sequence alignment (Kim, Paggi et al. 2019, Danecek, Bonfield et al. 2021). Quantification of sequencing data was achieved using featureCounts(Liao, Smyth et al. 2014). The genome fasta data and annotated gtf file were from NCBI(GCF_005281545.1), too. Scaled log 10 transformed counts was used for quality control checks in R, and 24299 genes were kept after qc filtration. Fig 1 C shows the RNA seq data was reliable for downstream analysis.

### 3. Sc-RNA seq

Raw data was processed with cellranger software. Then filtered and analyzed in R using Seurat (Hao, Hao et al. 2021, Hao, Stuart et al. 2023). Cells with nUMI more than 24000, nGene less than 400 and more than 5000, mitochondrial gene ratio more than 10% were omitted, 28698 cells with nGene median 1462 kept after qc. And reduce dimension using Seurat and integrate data with Harmony (Korsunsky, Fan et al. 2022),

fig 1 D shows the tSNE dimension reduction plot. The cell makers we used to identify cell types of groupers were described in the published literature we mentioned before (Wang, Peng et al. 2021). And we used cellchat for cell communication analysis next (Jin, Plikus et al. 2023), to overcome the difficulty that non-model organisms don’t have a stable ligand-receptor interaction relationship, we use STRINGdb to predict interactions in whole proteome level and built a cellchatdb of interested pathways to do cell communication analysis (Szklarczyk, Kirsch et al. 2023), we also developed an R package to simplify this procedure. Detailed analysis workflow was described in fig 1 A.

After data pre-processing, we first used limma to calculate statistical result on proteomic and transcriptomic data in R (Ritchie, Phipson et al. 2015). And then we focused on the intersection of differential expressed protein and genes. Luckily, the intersection result did show us some useful clues. Fig 2 shows the intersection of proteomic data and transcriptomic data. And table 1 shows the information of our interested genes and protein.

**Table 1:**
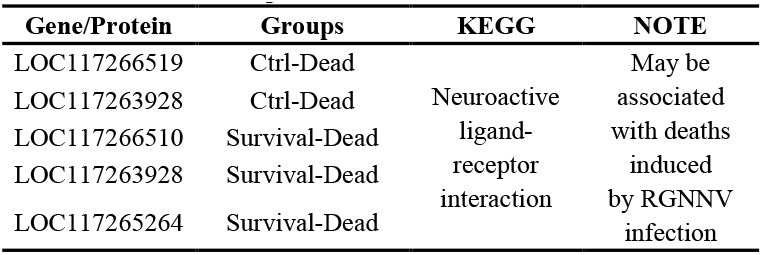
Important DEG/P list in intersection

**Fig. 2.**
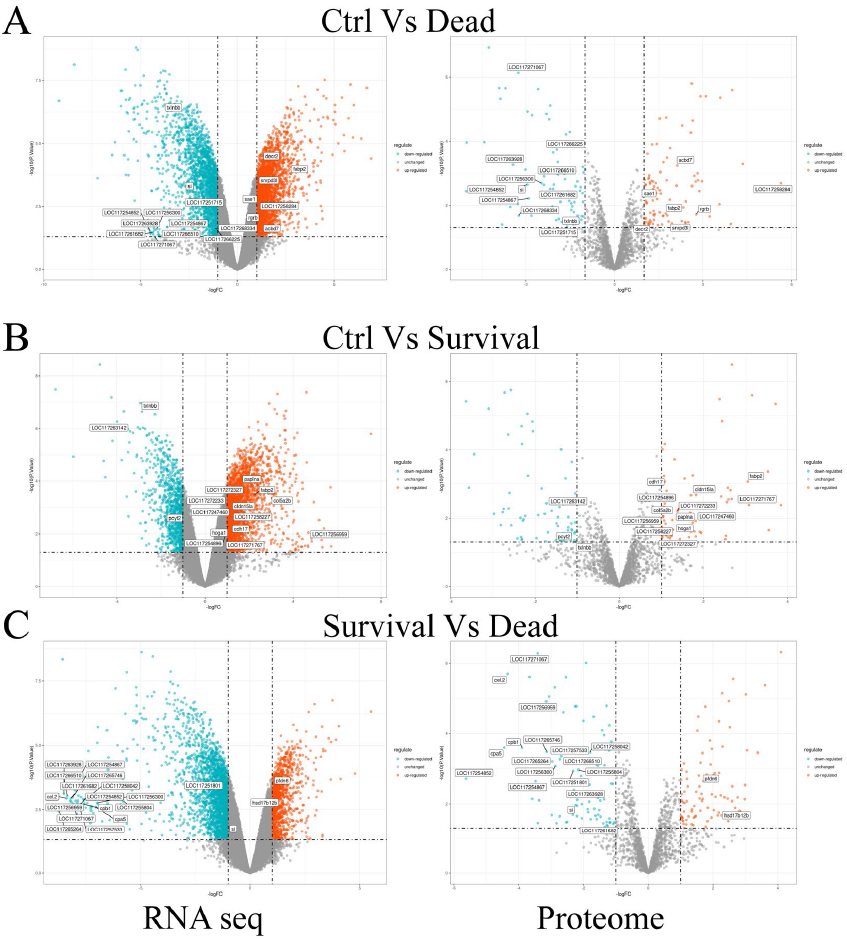
The intersection of proteomic data and transcriptomic data. A: Control Vs Infected/Dead. B: Control Vs Infected/Survival. C: Infected/Survival Vs Infected/Dead.

We noticed that after RGNNV infection, the neuroactive ligand-receptor interaction pathway (KEGG: ely04080) related genes were down regulated significantly, especially compared with the dead group. That may suggest that the RGNNV induced death may be due to the down regulate of the ely04080 pathway in grouper’s brain. To study the regulation mechanism of ely04080, we performed cell communication analysis with sc-RNA seq data.

The cell communication analysis would meet the difficulty that there’s no proper database in non-model organism. Normally, researchers do cell communication analysis by comparing homologous genes in humans or mice before. To overcome this difficulty, we developed a R package named ‘tocelldb’ which can be downloaded from github and first raise a solution to build a cellchatdb in non-model organisms those with a reference genome and a reference proteome.

We uploaded the reference proteome of *Epinephelus lanceolatus* to stringdb, then got the predicted interaction networks of our interested data from stringdb in tsv format. And we used tocelldb package to build a cellchatdb of our interested grouper’s pathways.

After getting the cellchatdb of neuroactive ligand-receptor interaction pathway, we performed cell communication with cellchat package in R. We found that glutamate receptor family is the key of how RGNNV disturb the normal metabolic process in grouper’s neuron system, and γ-aminobutyric acid (GABA) receptor family may be involved in this pathological process(fig 3 A).

**Fig. 3.**
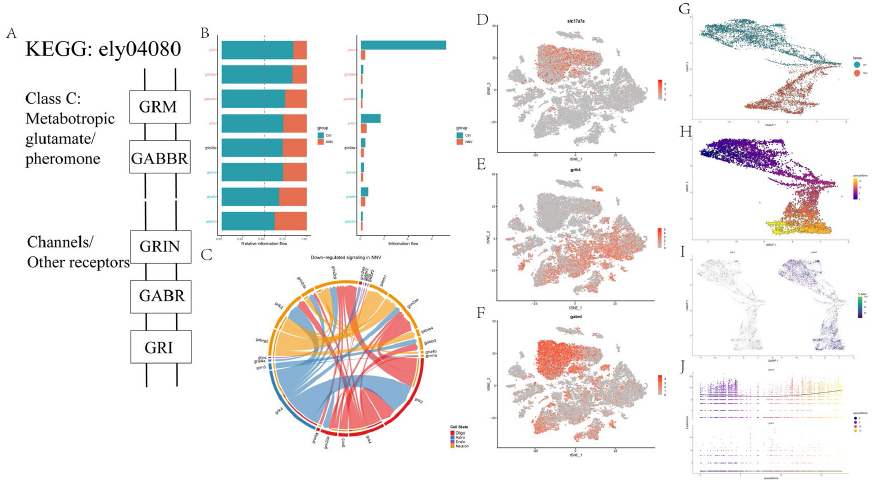
The inhibition of glutamate receptor family in neuroactive ligand-receptor interaction pathway. A: Inhibited pathways. B: Information flow of glutamate receptors and GABA receptors. B: Comparative information flow between infection and control groups. C: Down-regulated signaling in RGNNV infection. D: TSNE plot of slc17a7a, which reported as a gene susceptible to RGNNV attack. E, F: TSNE plot of grik4 and gabrd. G∼J: Cell trajectory of glu1 and glu3 subtypes. The consistent transcriptional level of grik4 throughout the infection progression suggests that glutamate receptor inhibition may not be induced by RGNNV infection but rather represents a specific genetic phenotype in the selected experimental animals.

Glutamate metabolism is an important part of normal nervous system activity and is usually responsible for mediating excitatory neurotransmitters. There’re four subfamilies of glutamate receptor, one of which is GRM (metabotropic glutamate receptor), mainly mediates the production of slow nerve signals. And the other three ionotropic glutamate receptor subtypes are GRIN (NMDA), GRIK (KAR) and GRIA (AMPA), respectively, mainly mediates fast nerve signal transmission.

GABA, in contrast to glutamate, is usually responsible for calming nerves and inhibiting excitability. The GABA receptor also divided in 2 subfamilies, the first one is GABBR, take charge of metabotropic pheromone function, And the later is GABR.

Normally, under the cooperative action of GABA and glutamate, neural activity in animals is generally maintained in a balanced state of excitation. But we noticed that after RGNNV infection, the activities of glutamate receptor family were significantly inhibited in neuron cells (fig 3 B, C).

This may suggest that, under RGNNV infection, there is a disruption in glutamate metabolism in the neural cells of grouper, induced by the inhibitory effects of glutamate receptors. Consequently, this disruption affects the normal neural activity in grouper, leading to abnormal neural excitation due to the accumulation of glutamate. This, in turn, triggers behavioral disorders in grouper.

In previously published literature, it has been demonstrated that the expression of glu1 and glu3 subtypes of slc17a7 is more susceptible to RGNNV attack (Wang, Peng et al. 2021). We are considering whether this susceptibility is related to glutamate receptor inhibition. The results indicate that indeed it is the case, especially grik4, and this characteristic is also associated with GABA receptors, gabrd, in particular, plays a crucial role in this process, too (fig 3 D, E, F).

To determine the transcriptional level changes of glutamate and GABA along a pseudotemporal trajectory, we performed single cell pseudotime analysis using Monocle 3 (Trapnell, Cacchiarelli et al. 2014, Qiu, Hill et al. 2017, Qiu, Mao et al. 2017, Cao, Spielmann et al. 2019). We differentiated GLU-type cells from GABA-type cells and conducted pseudotime analysis separately. The pseudotemporal trajectory clearly progressed towards RGNNV infection. However, we observed that the transcriptional levels of grik4 did not change throughout the entire infection timeline in the susceptible GLU-type cells (fig 3 G, H, I, J). This suggests that perhaps glutamate receptor inhibition is not induced by RGNNV infection but rather represents a specific genetic phenotype in the experimental animals. This indicates that if we identify characteristic genes regulated by glutamate receptors, it might be possible to screen fish fry with a certain level of immunity to RGNNV through genetic breeding, thereby reducing economic losses in grouper aquaculture caused by RGNNV infection to some extent.

These findings have directed our attention towards the glutamate metabolic pathway. While we understand that glutamate may be significantly associated with behavioral disorders and potential lethality in grouper, the mechanism through which glutamate metabolism disruption operates in this process remains unclear. Consequently, we have shifted our focus to the metabolic pathways involving alanine, aspartate, and glutamate (KEGG: ely00250). Once again, we are utilizing the predicted interaction data from stringdb to analyze changes in intercellular communication.

We are delighted to find that the clues have led us to the significant upregulation of glutamate dehydrogenase and glutamine synthetase in mitochondrial. Glutamine has previously been reported as a crucial amino acid for RGNNV replication, with its absence resulting in the inhibition of RGNNV replication without affecting the host’s metabolic activities.

In neural cells, glutamate is stored in vesicles at the presynaptic membrane through vesicular glutamate transporters. Upon depolarization of the presynaptic membrane, glutamate is released, binds to ionotropic glutamate receptors, and generates action potentials. Astrocytes regulate the release of excessive glutamate into the synaptic cleft through excitatory amino acid transporters against a concentration gradient. Within astrocytes, glutamate undergoes a cycle through the glutamine synthetase reaction, converting it to glutamine. The latter is then transported back to neurons and converted back to glutamate for use in new synapses (fig 4 A) (Silva, Souza et al. 2019). This indicates that through the interactive activities between neural cells and astrocytes, a glutamate-glutamine cycle is established, further exacerbating the accumulation of glutamate due to glutamate receptor inhibition. Simultaneously, we have observed increased communication activity in astrocytes following RGNNV infection in ely00250 pathway (fig 4 B), glulb serves as a crucial node in this cycle (fig 4 C, D).

**Fig. 4.**
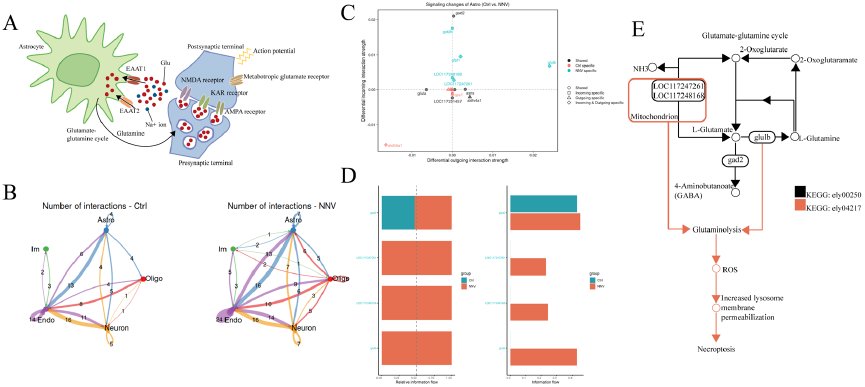
The necroptosis pathway (KEGG: ely04217) and alanine, aspartate, and glutamate metabolism pathway (KEGG: ely00250) may lead to death in grouper. A: The glutamate-glutamine cycle between astrocyte cells and neuron cells in healthy animal. B. Increased glutamate-glutamine cycle after RGNNV infection resulted in more active astrocytes. C. Signaling changes of astrocytes. D. Compare the overall information flow of ely00250. E. Diagram of regulatory patterns of necroptosis induced by increased glutamate-glutamine cycle. LOC117247261: glutamate dehydrogenase, mitochondrial, LOC117248168: glutamate dehydrogenase, mitochondrial-like, glulb: glutamine synthetase, gad2: glutamate decarboxylase 2 isoform X1.

Equally important, the glutamate-glutamine cycle is also a crucial component of the necroptosis pathway (KEGG: ely04217). After the formation of this cycle, it may induce an increase in reactive oxygen species (ROS), leading to lysosomal membrane permeabilization (LMP) and subsequent neuronal necrosis (fig 4 E).

Now, we have gained a comprehensive understanding of the mechanism by which RGNNV infection induces glutamate metabolism disruption, potentially leading to the death of grouper (fig 5). Glutamate metabolism is an important part of maintaining the stability of organism neural activity, and is related to the onset of Parkinson’s disease, Alzheimer’s disease, epilepsy and other neuron system diseases. The behavioral disorder of grouper infected with RGNNV may be like the pathogenesis of preprogrammed diseases. Understanding the mechanism by which RGNNV infects the midbrain system of grouper will not only help reduce the substantial economic losses of the grouper farming industry, but also help us understand the relationship between neural activity and glutamate metabolism in other animals.

**Fig. 5.**
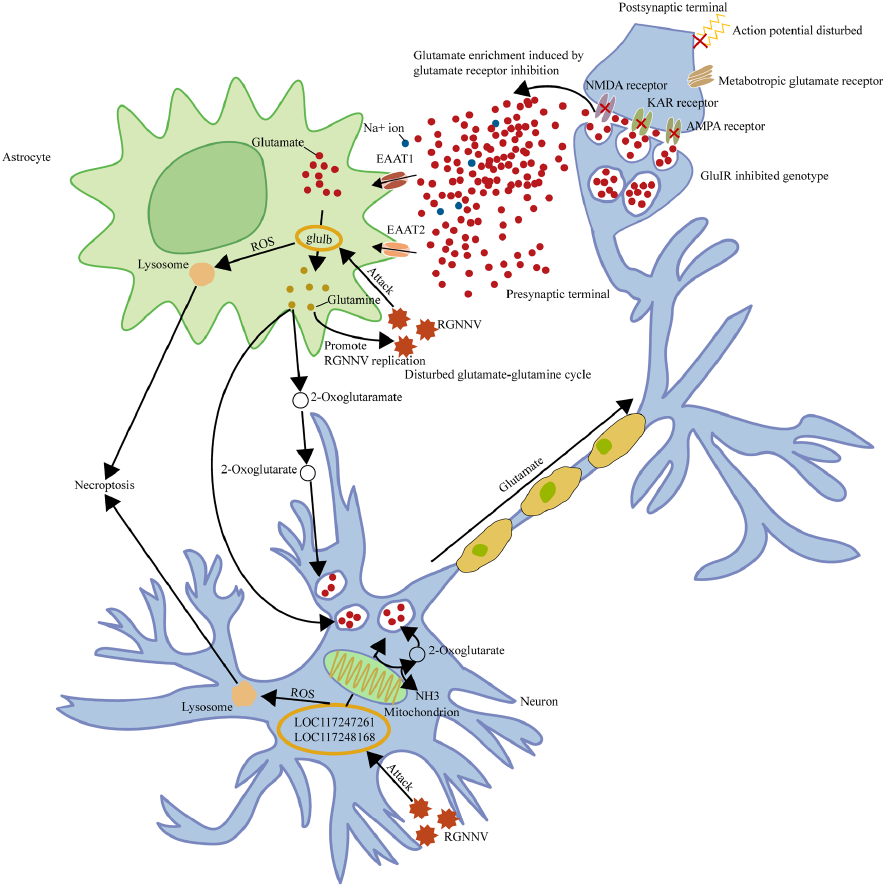
The schematic regulatory mechanism of glutamate accumulation induced by RGNNV infection in GluIR gene-inhibited grouper neural system. In GluIR gene-inhibited grouper neural systems, there is typically an accumulation of excess glutamate, leading to restricted action potentials and an active glutamate-glutamine cycle. Following RGNNV infection, this process is significantly upregulated. As RGNNV replication requires abundant glutamine, there is an initial activation of the glulb gene within astrocytes, producing more glutamine. The generated glutamine is then converted into new glutamate within neuronal cells through two pathways: one involves a direct biochemical reaction where glutamine is reduced to glutamate, and the other involves the transformation of intermediate products, 2-oxoglutaramate and 2-oxoglutarate, into glutamate. Additionally, within neuronal cells, RGNNV accelerates glutamate production by upregulating mitochondrial glutamate dehydrogenase, LOC117247261 and LOC117248168. The surplus glutamate is released through presynaptic terminals, but due to GluIR inhibition on postsynaptic bodies, a significant amount of glutamate is unable to pass through, leading to its accumulation at synapses. This excess glutamate is then transferred back to astrocytes, creating a vicious cycle. Furthermore, the heightened activity of glulb and mitochondrial glutamate dehydrogenase induces glutaminolysis, triggering the production of reactive oxygen species (ROS), altering lysosomal membrane permeability, and ultimately leading to necroptosis.

Additionally, our novel process for single-cell communication analysis in non-model organisms has been developed into an R package. This provides an analytical method for future single-cell communication analysis in all non-model organisms with reference. There are 2 main functions that provided by tocelldb package, the first is generating a cellchatdb with your interested genes/protein or a DEG/P list as input. The second is generating a cellchatdb with a predicted tsv file downloaded from stringdb of your interested KEGG/GO pathway or gene/protein (fig 6).

**Fig. 6.**
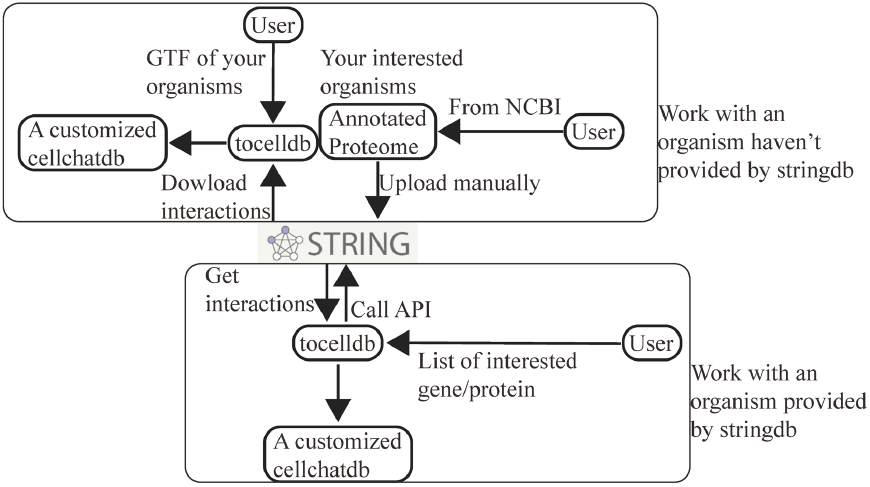
The diagram of functions provided in tocelldb package.

